# Surface-driven registration method for the structure-informed segmentation of diffusion MR images

**DOI:** 10.1101/018945

**Authors:** Oscar Esteban, Dominique Zosso, Alessandro Daducci, Meritxell Bach-Cuadra, María J. Ledesma-Carbayo, Jean-Philippe Thiran, Andres Santos

## Abstract

Current methods for processing diffusion MRI (dMRI) to map the connectivity of the human brain require precise delineations of anatomical structures. This requirement has been approached by either segmenting the data in native dMRI space or mapping the structural information from T1-weighted (T1w) images. The characteristic features of diffusion data in terms of signal-to-noise ratio, resolution, as well as the geometrical distortions caused by the inhomogeneity of magnetic susceptibility across tissues hinder both solutions. Unifying the two approaches, we propose *regseg*, a surface-to-volume nonlinear registration method that segments homogeneous regions within multivariate images by mapping a set of nested reference-surfaces. Accurate surfaces are extracted from a T1w image of the subject, using as target image the bivariate volume comprehending the fractional anisotropy (FA) and the apparent diffusion coefficient (ADC) maps derived from the dMRI dataset. We first verify the accuracy of *regseg* on a general context using digital phantoms distorted with synthetic and random deformations. Then we establish an evaluation framework using undistorted dMRI data from the Human Connectome Project (HCP) and realistic deformations derived from the inhomogeneity fieldmap corresponding to each subject. We analyze the performance of *regseg* computing the misregistration error of the surfaces estimated after being mapped with *regseg* onto 16 datasets from the HCP. The distribution of errors shows a 95% CI of 0.56–0.66 mm, that is below the dMRI resolution (1.25 mm, isotropic). Finally, we cross-compare the proposed tool against a nonlinear *b0*-to-T2w registration method, thereby obtaining a significantly lower misregistration error with *regseg*. The accurate mapping of structural information in dMRI space is fundamental to increase the reliability of network building in connectivity analyses, and to improve the performance of the emerging structure-informed techniques for dMRI data processing.

## 1. Introduction

Diffusion MRI enables the mapping of microstructure (Basser and Pierpaoli, 1996) and connectivity (Craddock et al., 2013) of the human brain *in-vivo*. It is generally acquired using echo-planar imaging (EPI) schemes, since they are very fast at scanning a large sequence of images called diffusion weighted images (DWIs). Each DWI is sensitized with a gradient to probe proton diffusion in a certain orientation. Subsequent processing involves describing the local microstructure with one of the available models, which range from the early diffusion tensor imaging (DTI) proposed by Basser and Pierpaoli (1996) to current models such as AMICO (accelerated microstructure imaging via convex optimization, Daducci et al., 2015). The microstructural map is then used to draw the preferential orientations of diffusion across the brain using tractography (Mori et al., 1999). Finally, a graph representing the corresponding structural network is built using the regions of a cortical parcellation as nodes and the fiber paths found by tractography as edges (Hagmann et al., 2008). The methodologies to solve reconstruction, tractography and network building require the delineation of the anatomy in the dMRI space. Moreover, current trends on reconstruction (Jeurissen et al., 2014) and tractography (Smith et al., 2012) are increasingly using structural information to improve the microstructural mapping and fiber-tracking.

Possibly, the earliest structural information incorporated to aid dMRI processing is the white matter (WM) mask used as a termination criteria for tractography. The standardized procedure to obtain this mask was thresholding the FA map. However, the mask and subsequent analyses are highly dependent on the threshold that is chosen (Taoka et al., 2009). To overcome the unreliability of FA thresholding, and to broaden WM segmentation to brain tissue segmentation, a large number of methods have been proposed using DWIs, the *b0*, and DTI-derived scalar maps such as FA, ADC and others (Zhukov et al., 2003; Rousson et al., 2004; Jonasson, 2005; Hadjiprocopis et al., 2005; Liu et al., 2007; Lu et al., 2008; Han et al., 2009). However, the precise segmentation of dMRI is difficult for several reasons. First, dMRI images have a resolution that is much lower than that of the imaged microstructural features. Therefore, voxels located in structural discontinuities are affected by partial voluming of the signal sources. Second, the extremely low signal-to-noise ratio (SNR) and the high dimensionality of the DWIs prevent their direct use in segmentation. Third, the low contrast between gray matter (GM) and WM in the *b0* volume also makes it unsuitable for brain tissue segmentation.

An alternative route to segmentation in dMRI space is the mapping of the structural information extracted from anatomical MR images, such as T1w, using image registration techniques. Generally, intra-subject registration of MR images of the brain involves only a linear mapping to compensate for head motion between scans. However, EPI introduces a geometrical distortion (Jezzard and Balaban, 1995) that impedes the linear mapping from the structural space. Numerous methods have been proposed to overcome this problem by incorporating information from extra MR acquisitions such as fieldmaps (Jezzard and Balaban, 1995), DWIs with a different phase-encoding (PE) scheme (Cordes et al., 2000; Chiou and Nalcioglu, 2000), or T2-weighted (T2w) images (Kybic et al., 2000; Studholme et al., 2000). These methods estimate the deformation field associated to *EPI distortions* and resample the DWIs onto a corrected dMRI space. The retrospective EPI correction is an active field of research yielding frequent refinements and combinations of the original methods, such as (Holland et al., 2010; Andersson et al., 2012; Irfanoglu et al., 2015). A standardized method to solve the remaining linear mapping between the corrected-dMRI and the structural spaces is *bbregister* (Greve and Fischl, 2009).

Here, we present a segmentation and surface-to-volume registration method called *regseg*, and show its usefulness in mapping anatomical information from structural space into native dMRI space to aid subsequent processing steps (reconstruction, tractography and network building using a cortical parcellation). The underlying hypothesis is that the registration and segmentation problems in dMRI can be solved simultaneously. To implement *regseg* we first establish an active-contours without edges (Chan and Vese, 2001) segmentation framework. A specific set of reference surfaces extracted from the same subject initialize the 3D active contours, which evolve searching for homogeneous regions in the multivariate target-image. We apply *regseg* to segment dMRI data by mapping a set of nested surfaces extracted from a structural image (e.g. T1w) to a bivariate target-volume comprehending the FA and ADC maps. The evolution of the surfaces is supported by a B-spline basis, optimized iteratively using a descent approach driven by shape-gradients (Jehan-Besson et al, 2003; Herbulot et al, 2006). Therefore, *regseg* establishes a registration framework that actually deals with the nonlinear warping induced by EPI distortions. *Regseg* integrates the benefits of segmentation and registration methods together and exploits the multivariate nature of dMRI data to contribute in the proposed application on three key aspects: 1) the surfaces are typically extracted from the T1w image of the same subject, therefore *regseg* does not require additional MR acquisitions to the minimal dMRI protocol in order to estimate the deformation field; 2) depending on the application, the resampling of the DWIs introduced by the correction method can be avoided, either by performing the posterior processing on the native dMRI space or applying the unwarping on the tractography itself; and 3) *regseg* increases the geometrical accuracy of the overall process. In this paper, we first verify the functionality of the method and the *regseg* implementation using a set of digital phantoms, demonstrating the sub-pixel accuracy in registration. Then, we evaluate *regseg* on real dMRI datasets, using a derivation of our instrumentation framework (Esteban et al, 2014a) which simulates known and realistic EPI distortions. We also compare *regseg* and a nonlinear registration method to map the *b0* to the corresponding T2w image of the same subject. This approach is the first step of the T2w-registration based (T2B) correction methods (Kybic et al, 2000). We reproduce the settings and implementation of a widely used diffusion processing software *(ExploreDTI*, Leemans et al, 2009). With this comparison, we demonstrate how *regseg* achieves higher accuracy with the simultaneous registration and segmentation process.

## 2. Methods

### 2.1. Registration framework and segmentation model

Let Γ*_R_* = {Γ*_m_*: *m* ∈ ℕ, *m* ≤ *N_S_*} be the set of *N_S_* surfaces extracted from the undistorted T1w image (the reference space *R*). We reformulate the segmentation of the distorted dMRI images (the moving space *M*) as a registration problem where we search for an underlying deformation field *U* such that the structures in *R* defined by Γ*_R_* align optimally with their corresponding structures in *M*:

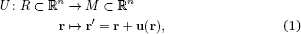

where r denotes a position in *R*, r’ is its corresponding location in *M*, and *n* denotes the dimensionality of images.

Finally, u = u(r) is the displacement of every point with respect to the reference domain. The general overview of how the surfaces interact with the registration framework is presented in Figure 1.

*Cost-function derivation*. In a Bayesian framework for registration (Wyatt and Noble, 2003; Pohl et al, 2006; Gass et al, 2014), the mappings *U* in (1) are evaluated based on their posterior probability given the observed data *M*. Let Ω = {Ω*_l_*: *l* ∈ ℕ, *l* ≤ *N_L_*} be the set of *N_L_* competing regions in which *M* is partitioned by the projection of Γ*_R_*. Using Bayes’ rule, the posterior likelihood is computed as:

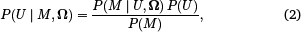

where *P*(*M* | *U*, *Ω*) is the data likelihood. Since Ω are mapped by *U*, we simplify *P*(*U*, Ω) = *P*(*U*) ⇒ *P*(*M* | *U*, Ω) = *P*(*M* | *U*). The best estimate *Û* then satisfies the maximum a posteriori criterion and it aligns Γ*_R_* into *M*. First, we assume independence between pixels, and thus we break down the global data likelihood into a product of pixel-wise conditional probabilities:

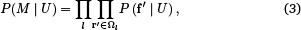

where f’ = *M*(r’) is the feature vector at the displaced position r’ (1) in the moving image. For convenience and because it has been shown to be an appropriate approximation (Van Leemput et al, 1999; Cuadra et al, 2005), we introduce two assumptions for each region Ω_*l*_: 1) the features are i.i.d.; and 2) they can be modeled by multivariate normal distributions *N*(f’ | *μ_l_*, Σ_*l*_;), with parameters {*μ_l_*, Σ_*l*_} for each region Ω_*l*_(Esteban et al, 2014b):

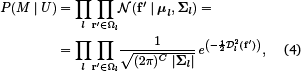

using 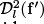 to denote the squared *Mahalanobis distance* of f’ with respect to the descriptors of region *l* as 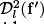 = (f’ – *μ_l_*)^T^ Σ_*l*_^−1^ (f’ – *μ_l_*). *C* is the number of channels comprised in the image *M*. Even though the features being segmented are not generally i.i.d., the spatial interdependency of voxels is implicitly supported by the piecewise smooth partition of the space Ω. In fact, the projection of Γ*_R_* onto *M* is an implicit segmentation model, for which the covariance matrix Σ_*l*_ of each region is minimized. The Figure 2 shows how the joint distribution of the input images is approximated with a mixture of multivariate normal distributions, and this minimization is illustrated for the segmentation of the FA and the ADC maps of one subject.

**Figure 1:**
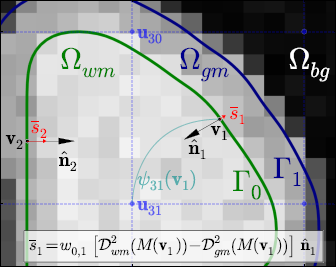
The interfacing surfaces Γ*_m_* between the competing ROIs Ω_*l*_, play the role of active contours which drive the registration process. They evolve iteratively along the normal 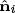 of the surface at each vertex v_*i*_ of the mesh. The gradient speeds 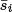 drive registration, which are computed as the disparity of the data energies with respect to the two limiting regions of *M*(v_*i*_), the features of the image *M* in the location of vertex v_*i*_. The computation of shape-gradients is developed in Appendix 2. In this figure, the 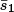 derived from Equation A.6 is written in the lower box, with Ω_*wm*_ in the inner limiting region, Ω_*gm*_ the outer region, and *w*_0,1_ is the relative area associated with vertex v_1_ with respect to the total area of surface Γ_0_.

*Regularization*. The smoothness of the resulting displacement field is induced by a Thikonov regularization prior:

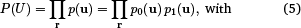

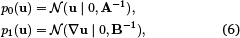

which requires that the distortion and its gradient have zero mean, and variance governed by the matrices **A** and **B**. Therefore, **A** and **B** are tensors that modulate the regularization, and produce deformations with preferential directions. Finally, the maximum a posteriori problem is adapted to a variational problem where we search for the minimum of an energy functional by applying *E*(u) = – log{*P*(*M* | *U*)*P*(*U*)}:

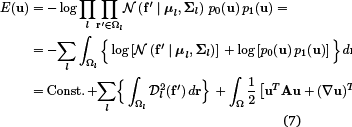

This expression is the dual of the Mumford-Shah functional that corresponds to the framework of active contours without edges (Chan and Vese, 2001) with the anisotropic regularization term of Nagel and Enkelmann (1986).

**Figure 2:**
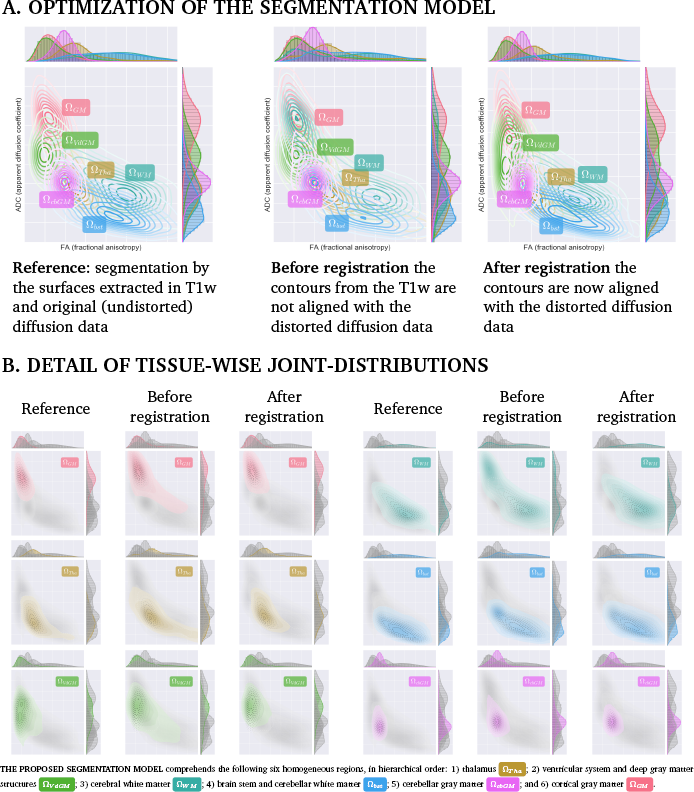
Evolution of the segmentation model defined by the homogeneous regions Ω_l_, for one real dataset. Panel (A, left) shows the joint distribution of the FA and ADC conditioned to the segmentation Ω defined by the surfaces **Γ***_R_* extracted from the T1w image. The plot was generated for reference using undistorted diffusion data, and therefore, **Γ***_R_* is aligned with the FA and the ADC. The problem arises when the diffusion data present deformation, and the contours **Γ***_R_* do not fit within the data (A, center). After registration with *regseg*, the contours are mapped onto the diffusion data (A, right), and the joint density plot is closer to the reference situation. In panel (B), the three plots in (A) are decomposed tissue-wise. Using filled contours, the bivariate distribution of each tissue is highlighted in its designated color, and represented over the remaining tissues (in gray colors). To help assessment, dashed contours in black-to-white colors represent the corresponding distribution in the reference plot. The registration process optimizes the segmentation model of *regseg*, and thus, the distribution of each region after registration is located closer to that corresponding in the reference situation, the shape of the distribution is more similar to the reference, and their spread is also reduced. The effects of optimization are more noticeable on the GM (Ω_GM_) and the WM (Ω_WM_). Particularly, the WM typ4ically shows a bimodal distribution when the contours **Γ** do not fit the data.

### 2.2. Numerical Implementation

*Deformation model*. Since the vertices of the surfaces {v*_i_*: v*_i_* ⊂ Γ}_*i*=1…*N_v_*_ are probably located off-grid, it is necessary to derive u*_i_* = u(v*_i_*) from a discrete set of parameters {u*_k_*}_*k*=1…*K*_. Densification is achieved using a set of associated basis functions *ψ_k_* (8). In our implementation, *ψ_k_* is a tensor-product B-spline kernel of degree three.

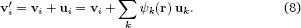

*Optimization*. To find the minimum of the energy functional (7), we propose a gradient-descent approach with respect to the underlying deformation field using the following partial differential equation (PDE):

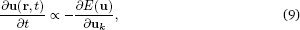

where *t* is an artificial time parameter of the contour evolution and u*_k_* are the parameters that support the estimate *Û* of the transformation at the current time point. Let us assume that the preferential directions of the displacement are aligned with the imaging axes to simplify (7) as expression (A.1) in Appendix 1, and thus to compute its derivative (9):

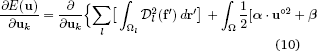

where u°^2^ = u^T^ · u, and {*α*,/*β*} are the expected variances along the imaging axes of the displacement field and its gradient, respectively. Then, the data and regularization terms are split and discretized to compute their derivatives. The derivative of the data term is computed using explicit shape gradients (see Appendix 2), which finally lead to obtain vertex-wise speeds of the gradient 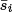 as illustrated in Figure 1. Using the expressions (A.7) and (A.8) given in Appendix 2 and introducing the analytical derivative of the regularization term, then the Equation 10 is reformulated as:

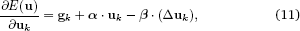

where g*_k_* are the shape-gradient contributions on the coefficients u*_k_* of the B-spline grid. Finally, to descend this gradient, we establish a semi-implicit Euler scheme (see Supplemental Materials, section S1.3), with a step size parameter *δ*, which we solve in the spectral domain as follows:

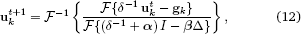

where *I* denotes the identity operator.

*Implementation details and convergence*. The *regseg* tool includes a multiresolution strategy on the free-form deformation field. Registration pyramids are created by setting the spacing between the control points of the B-spline basis functions for each level of the multiresolution strategy. As a rule of thumb, for a *δ* = 1.0, both *α* and *β* will typically be in the range [0.0,1.0]. The parameters used (*δ*, *α*, *β*, the B-spline grid resolutions, and target image smoothing), the implementation details, and other features such as the sparse matrix approach to fast interpolation are discussed in the Supplemental Materials, section S1.

### 2.3. Evaluation protocol

In order to assess the performance of *regseg*, we defined the following general evaluation protocol: 1) Extract the set of undistorted surfaces Γ*_R_*; 2) Compute a ground-truth field of displacements *U*_true_, which is applied to generate warped images (*M*) for segmentation; 3) Execute *regseg* with Γ*_R_* and use the warped data as inputs; and 4) Perform a visual assessment and compute the error metrics.

A first proof of concept is introduced to demonstrate *regseg* in digital phantoms with simple geometries, using *U*_true_ without directional restrictions. Then, *regseg* is evaluated in a framework using undistorted dMRI datasets, and u) t dr derived from the corresponding inhomogeneity fieldmap of the subject. Therefore, the deformation field is nonzero only in the phase-encoding (PE) axis, and reproduces a real EPI distortion. The adaptation of the evaluation protocol to the simulated phantoms and the real data is explained in the following sections.

### 2.4. Simulated phantoms

The workflow required to simulate the digital phantoms and to assess the performance of *regseg* with them is presented in Figure 3. A set of four binary objects (i.e. “box”, “ball”, “L”, and “gyrus”) was generated by combining the binarization of analytical shapes and mathematical morphology. The reference surfaces Γ*_R_* were extracted from the binary shapes using *FreeSurfer* tools (Fischl, 2012). The ground-truth distortion was generated using a chain of two displacement fields supported by grids of B-spline basis functions. The coefficients of the basis functions were generated randomly for both levels in their three dimensions. The three components of the displacements u = (*u_d_*) were bounded above by 40% of the separation between the control points at each level to obtain diffeomorphic transforms after concatenation (Rueckert et al, 2006). The first deformation field was applied to generate large warpings with control points separated by 50.50 mm in the three dimensions (*u_d_* ≤ 20.20 mm). With the second warping, we aimed to obtain a field with smoothness close to that found in a typical distortion field of dMRI data (Irfanoglu et al, 2011). Therefore, the control points were separated by 25.25 mm (*u_d_* ≤ 10.10 mm). After generating the ground-truth deformation, the original surfaces were warped by interpolating the displacement field at each vertex. The warped surfaces Γ_true_ were binarized to generate tissue fractions at low (2.0x2.0x2.0 [mm]) and high (1.0x1.0x1.0 [mm]) resolutions. Using an MRI simulator (Caruyer et al, 2014), we synthesized T1w (TE/TR= 10/1500 ms) and T2w images (TE/TR= 90/5000 ms), which corresponded to each phantom type, with each at two resolutions (1.0 mm and 2.0 mm isotropic). The field of view at both resolutions was 100x 100x [mm]. Next, *regseg* was applied to map Γ*_R_* onto the warped phantoms to obtain the registered surfaces (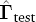). To quantify the misregistration error, we computed the Hausdorff distance between 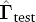 and Γ_true_ using (Commandeur et al, 2011). In total, 1200 experiments (four phantom types x 150 random warpings x two resolutions) were performed according to the workflow illustrated in Figure 3.

*Segmentation model and settings*. The segmentation model of the phantoms is implicitly defined: all phantoms comprehend an inner surface enclosing a uniform WM-like region, and an outer surface wrapping a GM-like layer. The outside is filled with uniform background (see Figure 3). All the experimental settings used for the phantoms are made available in a unique configuration file^1^.

### 2.5. Real datasets

The experimental framework for the real datasets is presented in Figure 4, which extends our previous evaluation (Esteban et al, 2014a) of distortions using dMRI phantoms.

*Data*. To evaluate *regseg* using real dMRI data obtained from human brains, we collected 16 subjects from the HCP database. The original acquisitions are released within “unprocessed” packages, whereas the “minimally preprocessed” packages contain the corresponding images after some processing (correction for several artifacts, brain-extraction, spatial normalization, etc.). We refer the reader to Van Essen et al. (2012) for exact details of the acquisition parameters and Glasser et al. (2013) for the preprocessing issues. These datasets comprise a large set of images, including T1w, T2w, and multi-shell dMRI images. Since we obtained the dMRI data from the minimally pre-processed package, these images are corrected for EPI distortions and spatially normalized in T1w space.

*Segmentation model*. Based on our experience and previous studies (Ennis and Kindlmann, 2006), we defined the moving image as a stack of the FA and ADC maps derived from dMRI data. After evaluating several alternative models, we empirically defined a partition *n* according to the following six regions: 1) thalamus (Ω*_Tha_*); 2) ventricular system and deep GM structures ( Ω*_V dGM_*); 3) cerebral WM (Ω*_WM_*); 4) brain stem and cerebellar WM (Ω*_bst_*); 5) cerebellar GM (Ω*_cbGM_*); and 6) cortical GM (Ω*_GM_*). Using tools in *FreeSurfer* and appropriate selections of labels in the *aparc* segmentation released with the HCP data, we extracted the Γ*_R_* set for the reference surfaces. The segmentation model corresponding to this partition is shown in Figure 2 and discussed in greater detail in the Supplemental Materials, section S4.

*Ground-truth generation*. Realistic deformation was achieved by generating displacement fields that satisfy the theoretical properties of distortion. The displacements along the PE axis of the dMRI image are related to the local deviation of the field Δ*B*0(r) from its nominal value *B*_0_ (Jezzard and Balaban, 1995), as follows:

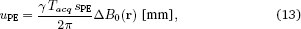

where *γ* is the gyromagnetic ratio, *T_acq_* is the readout time, and s _PE_ is the pixel size along PE. Certain MRI sequences are designed to estimate Δ*B*_0_, thereby obtaining the so-called fieldmap. We derived the deformation *U*_true_ from the fieldmap image released with the corresponding packages of each dataset in the HCP. The fieldmap was unwrapped^2^ and smoothed before applying (13). Next, the original dMRI was warped using the resulting displacement field and fed into a pipeline to process the corresponding DTI, thereby computing the derived scalars of interest (FA and ADC) using *MRtrix* (Tournier et al, 2012).

*Metric assessment*. Initially, we investigated the appropriateness of the segmentation model. For five test datasets, we uniformly sampled the space of distortions *Û* = *∊* · *U*_true_ = r + *∊ u*_PE_ (with *∊* ∈ [–1.1,1.1] and *u*_PE_ from (13)), and we evaluated the data term of the cost function (7). The minimum of the cost function (subsection 2.1) was consistently located at *∊* = 0.0 (the ground-truth) for all of the cases (Supplemental Materials, figure S2).

*Settings. Regseg* accepts an affine mapping from surface-space to the dMRI data as initialization. However, the images provided by the HCP are already spatially normalized. Therefore, the initial estimation of distortion is zero in this experiment. Since the distortion *U*_true_ is aligned along the PE direction (*y*-axis in our settings), *regseg* was configured to allow nonzero displacements only on that corresponding direction. For the experiments on real data, *regseg* established a multi-resolution pyramid of B-spline functions, with control points distributed on grids of the following spacings: 40x100x40 [mm] for the first (coarser) level, 30x30x30 [mm] for the second level, and 20x30x10 [mm] in the third level. Only the first and second levels included Gaussian smoothing of the target image (**σ** = [2.0, 0.5] mm, respectively). The actual choices of the parameter settings are publicly distributed with the source code for the experiments^3^. These settings were obtained manually, driven by the feedback obtained from the post-registration convergence reports (like that found in Supplemental Materials, section S1.3). We released *regseg* along with the tool to generate such convergence reports.

**Figure 3:**
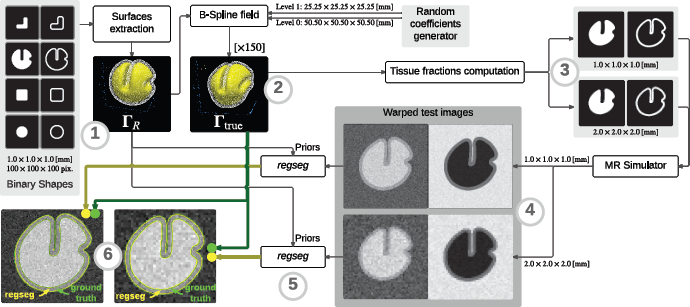
Evaluation of *regseg* using phantom data according to the following instrumental workflow. 1) The reference surfaces **Γ***_R_* are triangle meshes extracted from the four binary shapes (i.e., “box”, “ball”, “L”, “gyrus”). 2) A ground-truth displacement field was generated as described in subsection 2.4, and applied to warp **Γ***_R_*, thereby obtaining **Γ**_true_. 3) After being warped, Γ_true_ were projected onto the corresponding discrete 3D volume and downsampled to create partial volume effects at two resolutions, i.e., 2.0×2.0×2.0 [mm] and 1.0×1.0×1.0 [mm], thereby producing sets of tissue fractions maps. 4) The tissue fractions were fed into an MRI simulator, which generated T1-weighted (T1w) and T2-weighted (T2w) -like images at the two possible resolutions. 5) The *regseg* tool was applied using the warped test images as multispectral moving images and Γ_R_ as shape priors. 6) The agreement between the surfaces fitted by *regseg* (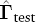) and **Γ**_true_ were assessed visually using automatically generated visual reports and quantitatively with the Hausdorff distance between the corresponding surfaces.

**Figure 4:**
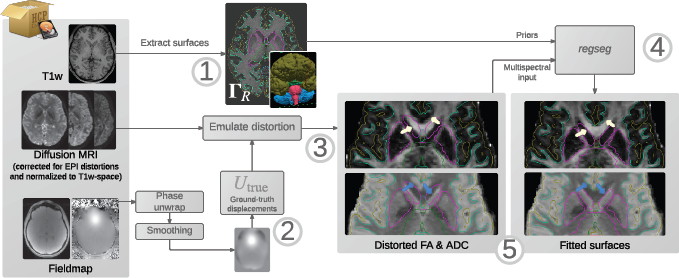
Experimental workflow employed to process real data from the Human Connectome Project (HCP). 1) **Γ***_R_* were extracted from the anatomical reference (T1w image). 2) For use as the ground truth, we generated a plausible synthetic distortion *U*_true_ from the fieldmap with (13). 3) The dMRI data were warped using *U*_true_ to reproduce the effects of real susceptibility-derived distortions. Target diffusion scalars (FA and ADC) were computed with the distorted data and stacked to feed the multivariate input required by *regseg*. 4) The method was run to obtain *U*_test_ = *Û*_true_, i.e., the estimate of the ground-truth deformation. 5) The results were evaluated visually and quantitatively. The arrows point to edges in the target images (light-yellow arrows for FA, blue for ADC) that should be aligned with a surface, showing how distortion limits the direct mapping from the structural space in which the contours are defined.

*Cross-comparison*. A dual workflow to the general evaluation used for *regseg* (Figure 4), was employed to integrate the alternate T2B registration scheme. We reproduced the solution and settings provided with *ExploreDTI* (Leemans et al, 2009), which is a widely used toolkit for tractography analysis of DTI. *ExploreDTI* internally employs *elastix* (Klein et al, 2010) to perform registration. The deformation field is correspondingly restricted to the PE direction. The settings file for *elastix* is also available^4^.

*Error measurement*. Distortion only occurs along the PE axis of the image, so we computed the surface warping index (sWI) as the area-weighted distance between the corresponding vertices of Γ_true_ and their estimate obtained by the method under the test 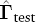:

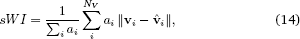

where **v**_*i*_ ⊂ Γ_true_ are the locations of the total *N_V_* vertices, *a_i_* is the area corresponding to each vertex **v***_i_*, and 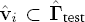 are the recovered locations that correspond to **v***_i_*. In practice, we only report the sWI for three surfaces ({Γ*_V dGM_*,Γ*_W M_*,Γ*_pial_*}) of crucial interest in whole-brain tractography.

## 3. Results

### 3.1. Verification and validation using digital phantoms

The results summarized in Figure 5 demonstrated that the accuracy was high and below the image resolution. Panel B on Figure 5 shows the violin plots for each model type corresponding to the two sets of resolutions for the generated phantoms. In order to relate the average misregistration error to the resolution of the moving image, we proceeded as follows. First, we confirmed that the

**Table 1:**
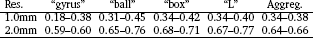
The distributions of vertex-wise Hausdorff distances between the ground-truth surfaces and their corresponding estimates obtained with *regseg* presented a 95% CI below the half-voxel size for all of the phantom types. The CIs were computed by bootstrapping using 10^4^ samples, with the median as the location statistic.

**Table 2:**
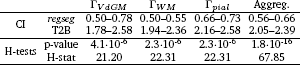
Statistical analysis of results obtained using 16 real datasets from the HCP, which show that *regseg* performed better than the alternative T2w-registration based (T2B) method. The distribution of the errors computed for the surfaces of interest (Γ*_V dGM_*, Γ*_W M_*, Γ*_pial_*) and the aggregate of all surfaces (Aggreg. column) had lower 95% CIs with *regseg*. The CIs in this table were computed by bootstrapping using the mean as the location statistic and with 10^4^ samples. The Kruskal-Wallis H-tests indicated that there was a significant difference between the results obtained using *regseg* and the T2B method.

vertex-wise error distributions were skewed by using Shapiro-Wilk’s test of normality. All of the distributions of errors in the tests (four phantom types × two resolutions) were nonnormal with *p* < 0.001. Consequently, we used the nonparametric Wilcoxon signed-rank test with the Bonferroni correction for multiple comparisons (*N*=150, for each phantom type). The average errors were significantly lower than the voxel size with *p* < (0.001/150) in all tests (four phantom types × two resolutions). Statistical tests might not be sufficiently conclusive, so we also computed the confidence intervals, as shown in Table 1.

### 3.2. Evaluation using real datasets and cross-comparison

Finally, we compared the performance of *regseg* with that of the standard T2B method. Summary reports for visual assessment of the 16 cases are included in the Supplemental Materials, section S5. In Figure 6, box A, the visual report is shown for one subject. We computed the sWI (14) of every surface after registration using both the *regseg* and T2B methods. Finally, to compare the results, we performed Kruskal-Wallis H-tests (a nonparametric alternative to ANOVA) on the warping indices for the three surfaces of interest selected in section 2.5 (Γ*_V dGM_*, Γ*_W M_*, Γ*_pial_*). All of the statistical tests showed that the error distributions obtained with *regseg* and T2B were significantly different, and the violin plots in box B of Figure 6 demonstrate that the errors were always larger with T2B. We also show the 95% CIs of the sWI for these surfaces (Table 2). The aggregate CI for *regseg* was 0.56-0.66 [mm], whereas the T2B method yielded an aggregate CI of 2.05-2.39 [mm]. The results of the statistical tests and the CIs are summarized in Table 2.

## 4. Discussion

We present *regseg*, a simultaneous segmentation and registration method that maps a set of nested surfaces into a multivariate target-image. The nonlinear registration process evolves driven by the fitness of the piecewise-smooth classification of voxels in the target volume imposed by the current mapping of the surfaces. We propose *regseg* to map anatomical information extracted from T1w images into the corresponding dMRI of the same subject. Previously, joint segmentation and registration has been applied successfully to other problems such as longitudinal object tracking (Paragios, 2003) and atlas-based segmentation (Gorthi et al., 2011). The most common approach involves optimizing a deformation model (registration) that supports the evolution of the active contours (segmentation), like Paragios (2003); Yezzi et al. (2003). *Regseg* can be seen as a particular case of atlas-based segmentation-registration methods, replacing the atlas by the structural image of the subject (*structure-informed segmentation*). The main difference of atlas-based segmentation and the application at hand is the resolution of the target image. Atlas-based segmentation is typically applied on structural and high-resolution images. A comprehensive review of joint segmentation and registration methods applied in atlas-based segmentation is found in (Gorthi et al., 2011). They also propose a multiphase level-set function initialized from a labeled atlas to implement the active contours that drive the atlas registration. Alternatively, *regseg* implements the active contours with a hierarchical set of explicit surfaces (triangular meshes) instead of the multiphase level sets, and registration is driven by shape-gradients (Herbulot et al., 2006). As an advantage, the use of explicit surfaces enables segmenting dMRI images with accuracy below pixel size.

**Figure 5:**
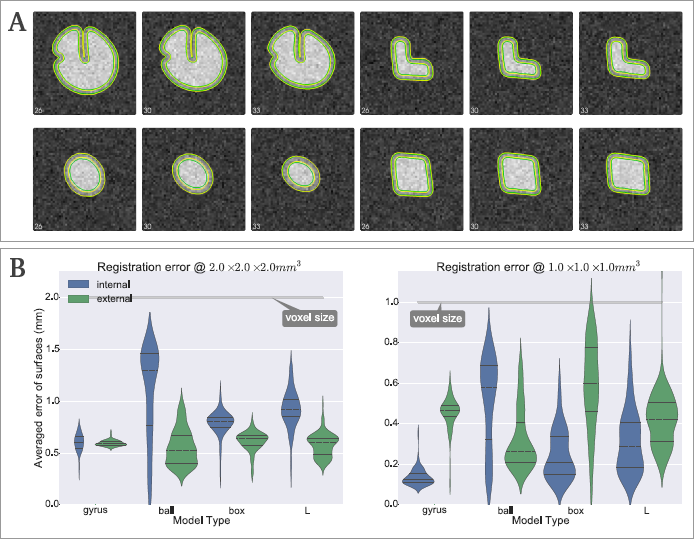
A. Visual assessment of the results obtained on the digital phantoms. The panel shows three coronal slices (indices indicated on the lower left corner) of each phantom volume at low resolution: “gyrus” (top left), “L” (top right), “ball” (bottom left), and “box” at (bottom right). The contours recovered after registration are represented in yellow. *Regseg* achieved high accuracy because it determined the almost exact locations of the registered contours with respect to their ground truth positions (shown in green). The partial volume effect makes segmentation of the sulci a challenging problem with voxel-wise clustering methods, but they were successfully segmented with *regseg*. B. Quantitative evaluation: the violin plot shows the variability across experiments of the average Hausdorff distance measured in each vertex of the corresponding surface, for the low (left) and high (right) resolutions. Error averages were consistently below the size of the voxel.

**Figure 6:**
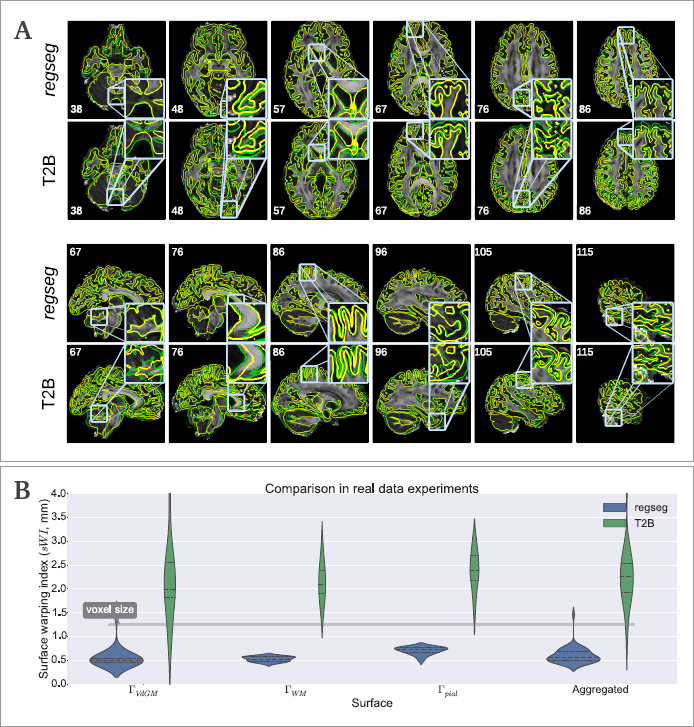
A. Example of a visual assessment report, which was automatically generated by the evaluation tool. Each view shows one component of the input image (in this case, the FA map), the ground-truth locations of the surfaces (green contours), and the resulting surfaces obtained with the test method (yellow contours). The first two rows show axial slices for *regseg* and the T2w-registration based (T2B) method, while the last two rows show the corresponding sagittal views. The coronal view is omitted because it was the least informative due to the directional property of the distortions. Specific regions where *regseg* outperformed T2B are enlarged. B. Violin plots of the error distributions for each surface across the 16 subjects, which show the voxel size of the dMRI images (1.25 mm), thereby supporting the improved results obtained by *regseg* with the proposed settings.

An important antecedent of *regseg* is *bbregister* (Greve and Fischl, 2009). The tool has been widely adopted as the standard registration method to be used along with the EPI correction of choice. It implements a linear mapping and uses 3D active contours *with edges* to search for intensity boundaries in the *b0* image. The active contours are initialized using surfaces extracted from the T1w using *FreeSurfer* (Fischl, 2012). To overcome the problem of nonlinear distortions, *bbregister* excludes from the boundary search those regions that are typically warped. Indeed, the distortion must be addressed separately because it is not supported by the affine transformation model. Conversely, the deformation model of *regseg* is nonlinear and the active contours are *without edges* (Chan and Vese, 2001) since the FA and ADC maps do not present steep image gradients (edges) but the anatomy can be identified by looking for piece-wise smooth homogeneous regions.

Recently, Le Guyader and Vese (2011) proposed a simultaneous segmentation and registration method in 2D using level sets and a nonlinear elasticity smoother on the displacement vector field, which preserves the topology even with very large deformations. *Regseg* includes an anisotropic regularizer for the displacement field proposed by Nagel and Enkelmann (1986). This regularization strategy conceptually falls in the midway between the Gaussian smoothing generally included in most of the existing methodologies, and the complexity of the elasticity smoother of Le Guyader and Vese (2011). Other minor features that differ from current methods in joint segmentation and registration are the support of multivariate target-images and the efficient computation of the shape-gradients implemented with sparse matrices.

We verified that precise segmentation and registration of a set of surfaces into multivariate data is possible on digital phantoms. We randomly deformed four different phantom models to mimic three homogeneous regions (WM, GM, and cerebrospinal fluid) and we used them to simulate T1w and T2w images at two resolution levels. We measured the Hausdorff distance between the contours projected using the ground-truth warping and the estimations found with *regseg*. We concluded that the errors were significantly lower than the voxel size. We also assessed the 95% confidence interval (CI), which yielded an aggregate interval of 0.64–0.66 [mm] for the low resolution phantoms (2.0 mm isotropic voxel) and 0.34–0.38 [mm] for the high resolution phantoms (1.0 mm isotropic). Therefore, the error was bounded above by half of the voxel size. The distributions of errors along surfaces varied importantly depending on the shape of the phantom (see Figure 5B). The misregistration error of the “gyrus” phantom showed a much lower spread than that for the other shapes. We argue that the symmetry of those other shapes posed difficulties in driving the contours towards the appropriate region due to *sliding* displacements between the surfaces and their ground-truth position. The effect is not detectable by the active contours framework, but it is controllable increasing the regularization constraints. When *regseg* is applied on real datasets, this surface sliding is negligible for the convoluted nature of cortical surfaces and the directional restriction of the distortion.

We evaluated *regseg* in a real environment using the experimental framework presented in Figure 4. We processed 16 subjects from the HCP database using both *regseg* and an in-house replication of the T2w-registration based (T2B) method. *Regseg* obtained a high accuracy, with an aggregate 95% CI of 0.56–0.66 [mm], which was below the pixel size of 1.25 mm. The misregistration error that remained after *regseg* was significantly lower (*p* < 0.01) than the error corresponding to the T2B correction according to Kruskal-Wallis H-tests (Table 2). Visual inspections of all the results (Supplemental Materials, section S5) and the violin plots in Figure 6 confirmed that *regseg* achieved higher accuracy than the T2B method in our settings. We carefully configured the T2B method using the same algorithm and the same settings employed in a widely-used tool for dMRI processing. However, cross-comparison experiments are prone to the so-called *instrumentation bias* (Tustison et al., 2013). Therefore, these results did not prove that *regseg is better than* T2B, but indicated that *regseg* is a reliable option in this application field. Finally, we also proposed a piecewise-smooth segmentation model defined by a selection of nested surfaces to partition the multispectral space comprehending the FA and the ADC maps and ultimately identify anatomical structures in dMRI space. We also demonstrated the smoothness of the objective function on five of the real datasets (Supplemental Materials, figure S2), taking advantage of the directional restriction of possible distortions. However, *regseg* requires densely sampled surfaces to ensure the convergence. Using the digital phantoms, we severely decimated the surfaces by a large factor. These surfaces introduced a bias which displaced the zero of the gradients from the minimum of the objective function impeding the convergence.

The proposed application of the method in the task of identifying structural information in dMRI images is an active field of research (Jeurissen et al., 2015). Current processing of dMRI involved in the connectome extraction and other applications (such as tract-based spatial statistics (TBSS) or surgical planning) require a precise segmentation of the anatomical structures in the diffusion space. Some examples of these processing tasks are the structure-informed reconstruction of dMRI data (Jeurissen et al., 2014; Daducci et al., 2015), the anatomically constrained tractography (Smith et al., 2012), and the imposition of the cortical parcellation mapped from the T1w image (Hagmann et al., 2008). The problem was firstly addressed using image segmentation approaches in the native diffusion space, without definite and compelling results. With the introduction of retrospective correction methods for the *EPI distortions* and image registration approaches, the task has been typically solved in a two-step approach. First, the DWIs are corrected for *EPI distortions* by estimating the nonlinear deformation field from extra MR acquisitions (Jezzard and Balaban, 1995; Chiou and Nalcioglu, 2000; Cordes et al., 2000; Kybic et al., 2000). Second, mapping the structural information from the corresponding T1w image using a linear registration tool like *bbregister* (Greve and Fischl, 2009). The current activity on improving correction methods (Irfanoglu et al., 2015) and the comeback of segmentation of dMRI in its native space (Jeurissen et al., 2015) proof the open interest of this application. *Regseg* addresses this joint problem in a single step and it does not require any additional acquisition other than the minimal protocol comprehending only T1w and dMRI images. This situation is found commonly in historical datasets. We envision *regseg* to be integrated in diffusion processing pipelines, after a preliminar DTI computation and before anatomically-informed reconstruction and tractography methods. Since the structural information is projected into the native space of dMRI, these two processes and the matrix building task can be performed on the unaltered dMRI signal (i.e. without resampling data to an undistorted space). For analyses other than connectivity, like TBSS, the deformation estimated by *regseg* can be used to map the tracts into structural space. Beyond the presented application on dMRI data, *regseg* can be indicated in situations where there are precise surfaces delineating the structure, a target multivariate image in which the surfaces must be fitted, and the mapping between the surfaces and the volume encodes relevant physiological information, such as the normal/abnormal development or the macroscopic dynamics of organs and tissues. For instance, *regseg* may be applied in fields like neonatal brain image segmentation in longitudinal MRI studies of the early developmental patterns (Shi et al., 2010). In these studies, the surfaces obtained in a mature timepoint of the brain are retrospectively propagated to the initial timepoints, regardless of the changes in the contrast and spatial development between them. More generally, *regseg* may also be applied to the personalized study of longitudinal alteration of the brain using multi-spectral images, for instance in the case of traumatic brain injury (Irimia et al., 2014) or in monitoring brain tumors (Weizman et al., 2014).

## Conclusion

*Regseg* is a variational framework for the simultaneous segmentation and registration of 3D dMRI data obtained from the human brain, where within-subject anatomical information is used as a reference. The registration method segments the target multivariate image into several competing regions, which are defined explicitly by their limiting surfaces. The surfaces are active and they evolve on a free-form deformation field supported by the B-spline basis. A descent optimization strategy is guided by shape gradients computed on the current partition of the target image. *Regseg* uses active contours without edges and it searches for homogeneous regions within the image. We tested *regseg* using digital phantoms by simulating T1w and T2w MRI warped with smooth and random deformations. The resulting misregistration of the contours was significantly lower than the image resolution of the phantoms.

We proposed *regseg* for simultaneously segmenting and registering dMRI data to their corresponding T1w image from the same subject. We demonstrated the accuracy of the proposed method based on visual assessments of the results obtained by *regseg* and cross-comparisons with a widely used technique. Moreover, *regseg* does not require any images in addition to the minimal acquisition protocol, which only utilizes T1w and dMRI. As well as the proposed application to dMRI data, other potential uses of *regseg* are atlas-based segmentation and tracking objects in time-series.

## Availability and reproducibility statement

We considered the reproducibility of our results as a design requirement. Therefore, we used real data obtained from the Human Connectome Project (Van Essen et al., 2012) and all of the software utilized in this study is also publicly available. *Regseg* was implemented on top of ITK-4.6 (Insight Registration and Segmentation Toolkit, http://www.itk.org). The evaluation instruments (Figure 4) were implemented using *nipype* (Gorgolewski et al., 2011) to assess their reproducibility. All of the research elements (data, source code, figures, manuscript sources, etc.) involved in this study are publicly available under a unique package (Esteban and Zosso, 2015).

## Author contributions

All the authors contributed to this study. OE implemented the method, designed and conducted the experiments, wrote the paper, simulated the phantoms, and prepared the real data. DZ devised and drafted the registration method, generated early phantom datasets, and collaborated in the implementation of the method. AD, MBC, and MJLC interpreted the results. AD, MBC, MJLC, JPT, and AS advised on all aspects of the study.

## Appendix

*Appendix 1. Simplifying the regularization term*

The exponentials of the Thikonov regularization prior (6) have the general form v*^T^*Mv. If M is a *n* × *n* diagonal matrix such that M = mI*_n_*, then:

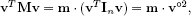

where we have introduced the Hadamard power notation^5^.

In general, the anisotropy of the distortion field is aligned with the voxel coordinate system, so A and B of (7) can be simplified to diagonal matrices to regularize the registration process, such that A = *α* I_*n*_ and B = *β*I_*n*_. By substituting into equation (7), we obtain:

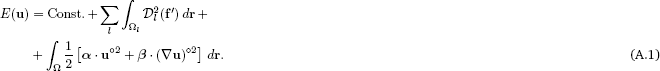

*Appendix 2. Application of the shape-gradients*

The computation of gradients at the locations of the active contours in the instant t is based on the work of Herbulot et al. (2006). Let F(r) be an “arbitrary” function over the image domain Ω = Ω_*l*_ ∪ Ω*_m_* split in two regions *l* and *m*, and Γ*_l,m_* a closed boundary between them. We now derive the domain integral w.r.t. *t*:

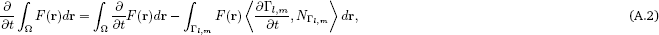

where 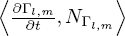 is the projection of the boundary movement on the unit inward normal *N*_Γ*_l,m_*_. Assuming that the region descriptors {*μ_l_*, Σ_*l*_} vary slowly enough, we can consider that 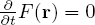 and thus:

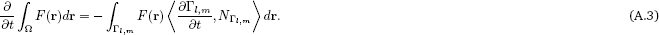

The equation (A.3) is discretized as follows. First, the surface between limiting regions *l* and *m* ( Γ*_l,m_*) is explicitly represented by a discrete set of vertices v*_i_*, with *i* ∈ {0,…,*N_p_* – 1}. Consequently, the inwards normal of the surface *N*_Γ*_l,m_*_ is represented by the discrete set of normals 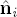 at each vertex of the mesh. The resulting summation is, therefore, discrete and the integral operator is replaced by the sum:

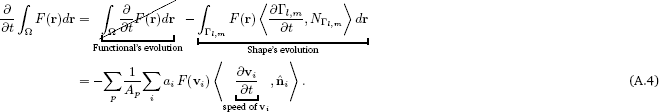

where *a_i_* is the area corresponding to vertex v*_i_*, and *A_p_* = £_*i*_ *a_i_* is the total area of surface *p*. In the following, we will refer as *w_p,i_* = *a_i_*/*A_p_* to the area contribution of v_*i*_ to the total area of the surface it belongs to. For simplicity, the sum over *p* can be also removed, as the vertices belong to only one of the total *P* contours.

Then, the speed of v_*i*_ is discretized using the artificial time-step parameter *δ*, as the displacement 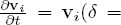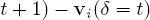

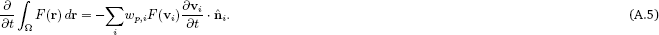

Since the energy functional is defined over competing regions, the displacement of v_*i*_ will cause an energy exchange between the limiting regions, and therefore *F*(*r*) must be split in two terms, *F_in_*(*r*) corresponding to the interior region and *F_out_*(*r*) to the exterior:

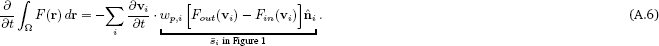

Then, we identify the shape gradient contribution g*_k_* on the coefficients u*_k_* of the B-spline grid:

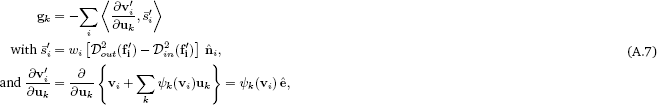

where *ê* is the coordinates system’s unit vector. Therefore, the shape gradients projected to the grid of B-spline control points read:

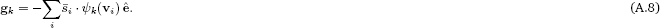

## Footnotes

* Supplemental materials to this work have been submitted to Data in Brief (Esteban et al., 2015). During the review process, the document is available in doi: 10.6084/m9.figshare.1397502.

1 https://github.com/oesteban/RegSeg/blob/master/Scripts/pyacwereg/data/regseg_default.json

2 fieldmaps are phase maps, which are intrinsically clipped in the interval of [-π, π) [rads] or [rads/s].

3 https://github.com/oesteban/RegSeg/tree/master/Scripts/pyacwereg/data/regseg_hcp.json

4 https://github.com/oesteban/RegSeg/blob/master/Scripts/pyacwereg/data/t2b_elastix_y.txt

5 The Hadamard power of a matrix or a vector is the power of its elements M^°*p*^ = (*m_ij_^p^*)

## Acknowledgments

The authors thank Y. Alemán and G. Wollny for their thorough reviews of the manuscript, V. Estellers for early discussions at the beginning of this project, and L. A. Vese for her support during OE’s research visits to her laboratory. We also thank A. Leemans for kindly sharing a p-code version of *ExploreDTI* from which the settings for *elastix* could be extracted.

DZ was supported by the Swiss National Science Foundation under grants PBELP2-137727, P300P2-147778, and NSF-DMS 1418812. This study was supported by the Spanish Ministry of Science and Innovation (projects TEC-2013-48251-C2-2-R and INNPACTO XIORT), Comunidad de Madrid (TOPUS) and European Regional Development Funds, the Center for Biomedical Imaging (CIBM) of the Geneva and Lausanne Universities and the EPFL, as well as the Leenaards and Louis Jeantet Foundations.

